# Immunostimulatory guide RNAs mediate potent antiviral response

**DOI:** 10.1101/282558

**Authors:** Yujia Cai, Alice Knudsen, Samuel Joseph Windross, Martin K Thomsen, Søren R Paludan

## Abstract

Genome-editing with CRISPR has emerged as a technology with broad therapeutic potential. However, it is unclear whether CRISPR will elicit innate immune responses, which could impact both positively and negatively on the desired therapeutic effects. Here, we have examined the immune-stimulatory properties of different variants of guide RNAs (gRNAs) – *in vitro* transcribed gRNA (IVT-gRNA) and synthetic gRNAs with or without chemical modifications, full-length or duplexed. We find that only IVT-gRNA evokes strong expression of cytokines in a panel of cell lines while all the synthetic RNAs do not. We further find that sensing of IVT-gRNA proceeds mainly through the RIG-I/MAVS RNA sensing axis. One potential use of CRISPR is for antiviral therapy. The antiviral actions of the gRNA tested up until now have been relying purely on the gene editing function of the CRISPR machinery, which weakens its feasibility due to the difficulty to target all infected cells. When IVT-gRNA was combined with unmodified Cas9 mRNA, which also induces cytokine expression, strong immune response was obtained while maintaining nuclease activity of CRISPR. Remarkably, such combination inhibited herpes simplex virus type-1 (HSV-1) replication even though the nuclease activity was modest, and provided ‘bystander protection’ to the cells that were not transfected with CRIPSR molecules. The antiviral activity of IVT-gRNA was also observed *in vivo* in HSV-1-infected Cas9+ mice, thus demonstrating the therapeutic potential. Our study further extends the applications of CRISPR by exploiting the immunostimulatory function of gRNAs.

## Introduction

CRISPR is now broadly used in basic and translational research. The technology was originally invented for its site-specific nuclease activity (Jinek et al. 2012; Cong et al. 2013; Jinek et al. 2013), and has subsequently been broadened to also allow transcription activation, nucleotide changing, and epigenetic reprograming (Maeder et al. 2013; Mali et al. 2013; Perez-Pinera et al. 2013; Komor et al. 2016; Vojta et al. 2016). For a successful clinical translation, understanding the host response to CRISPR is crucial as the potential immune reaction could conceivably lead to compromised gene editing efficiency and clearance of modified cells. It has been noted that long-term Cas9 exposure elicits Cas9 specific T-cell expansion and antibody production in mice (Chew et al. 2016). Prior to adaptive immune responses to Cas9 protein, the CRISPR can be sensed by the innate immune system which can detect foreign nucleic acids and is normally involved in recognition of invading viruses and bacteria. So far, it is unclear if gRNA - another component of CRISPR - will be recognized by the targeted cells. The gRNA which guides Cas9/gRNA complex to an intended genomic locus is a ∼100-nucleotides sequence consisting of both single- and double-stranded structures. It can be transcribed intracellularly driven by U6 promoter or can be delivered directly in the RNA format from *in vitro* transcription or synthesis. The *in vitro* transcribed gRNA (IVT-gRNA) together with Cas9 mRNA have been commonly used for microinjection into one-cell-stage embryos to quickly generate gene-modified animals (Niu et al. 2014). The synthetic gRNA with chemical modifications has shown superior efficiency and demonstrates potential in gene therapy research (Hendel et al. 2015; Dever et al. 2016). Compared to the well characterized nuclease activity, it is unclear if administration of gRNAs causes additional effects in the intracellular environment such as innate immune response.

In this study, we investigated the innate immune stimulatory properties of gRNAs from different sources – *in vitro* transcribed gRNA and a variety of synthetic gRNAs. In several cell types, we found that IVT-gRNA, but none of the synthetic gRNAs, elicits strong expression of a panel of cytokines including *IFNB1, TNFA, IL6* and *CXCL10* while maintaining nuclease activity. Taking advantage of this dual function of IVT-gRNAs, we demonstrated that they induced highly efficient antiviral response and provided bystander protection to the neighbouring non-transfected cells. By using a HSV-1 infection mouse model, we further extend the robust antiviral effects of IVT-gRNA *in vivo*.

## Results

### *In vitro* transcribed - but not the synthetic gRNAs - induce strong cytokine responses

Cells express a variety of pattern recognition receptors (PRRs), which sense danger signals through binding to microbial nucleic acids, proteins and carbohydrates (Mogensen 2009). The recognition of external substrates by PRRs initiates a hierarchical innate immune signalling leading to activation of specific gene expression programmes. In particular, intracellular nucleic-acid-sensing PRRs induce expression of type I and type III IFNs, which have strong antiviral activity (Mogensen 2009). To characterize the immunostimulatory properties of gRNAs, we first transfected THP1-derived macrophage-like cells with five types of gRNA targeting to the endogenous *CCR5* locus. The gRNAs were *in vitro* transcribed gRNAs (IVT) and four synthetic variants of gRNAs, namely full-length unmodified (FL-UM) and modified (FL-MO) gRNAs, and duplexed unmodified (Du-UM) and modified (Du-MO) gRNAs. We then examined innate immune response by measuring expression of the cytokines, *IFNB1, TNFA, CCL5, IL6*, and *CXCL10* as well-as the IFN-stimulated gene *MX1*. We found that IVT-gRNA induced robust cytokines expression to an extent comparable to the double-stranded RNA mimic poly(I:C), which is a known potent inducer of innate immune responses (Fig. 1A-F). The strong induction of *IFNB1* expression by IVT-gRNA was also seen in HEK293T cells and the human plasmacytoid dendritic cell line pDC-gen2.2 (Fig. 1G-H). Notably, none of the synthetic gRNAs enhanced the expression of the genes examined – regardless the chemical modifications (Fig. 1A-H). The immune stealth property of synthetic gRNAs might make them a preferred choice for *ex vivo* gene therapy where limited immune activation is desired and to preserve the ‘stemness’ of hematopoietic stem cells (Piras et al. 2017; Takizawa et al. 2017). To test if innate immune activation by gRNAs was dependent on transfection reagent, we transfected gRNAs with 3 different cationic lipid or polymer reagents. IVT-gRNA preserved the ability to induce *IFNB1* expression while the modified synthetic gRNA did not evoke any significant cytokine expression with all the transfection reagents (Fig. 1I and Supplemental_Fig_S1A). As CRISPR nuclease function requires formation of gRNA and Cas9 complexes, we also evaluated the commonly used Cas9 sources – the modified and unmodified Cas9 mRNA. While unmodified Cas9 mRNA clearly induced expression of cellular expression of immune genes, notably *IFNB1* and *TNFA*, the modified Cas9 mRNA was immune silent (Supplemental_Fig_S2A-F).

**Fig. 1.**
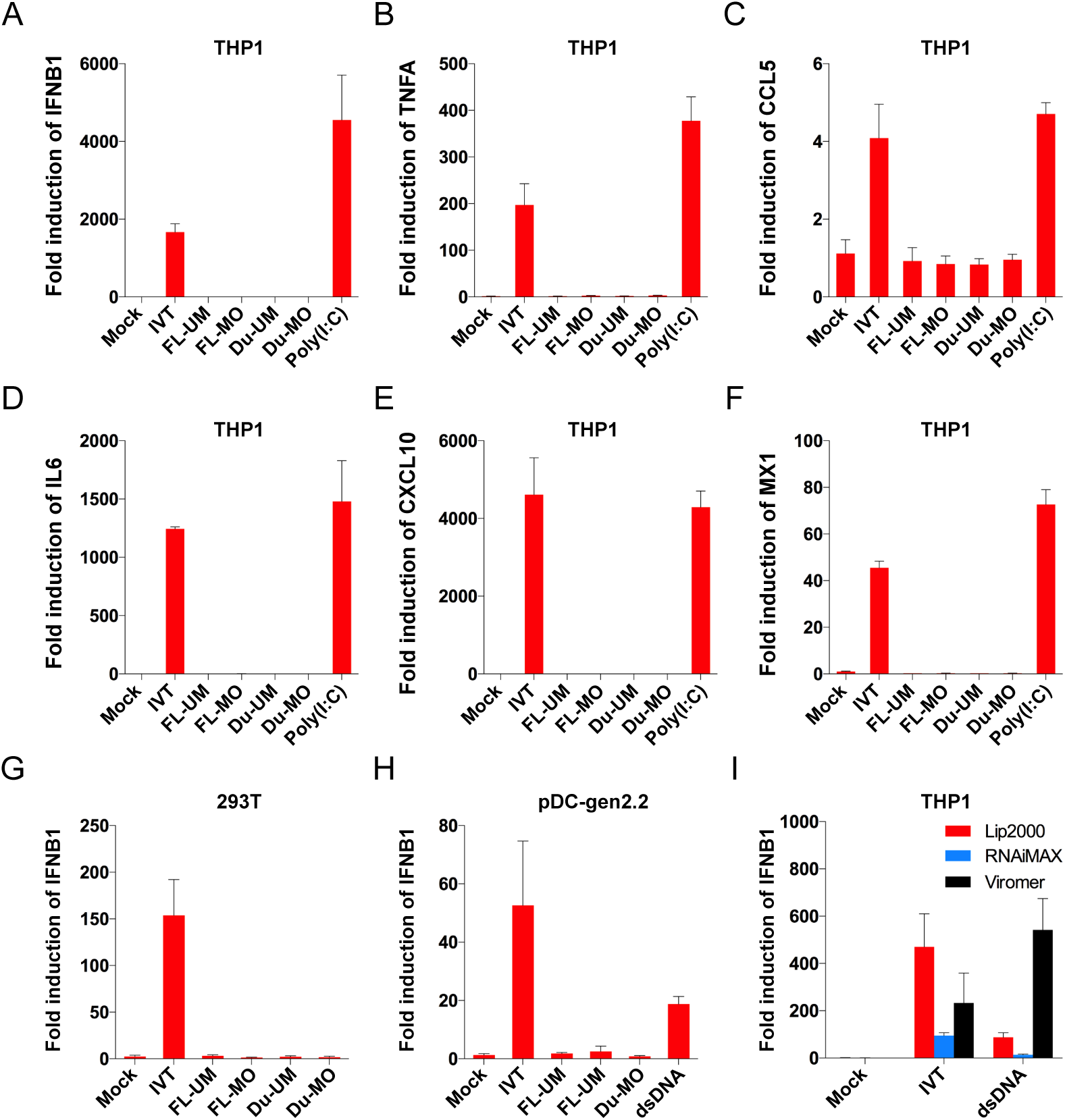
The innate immune property of gRNA variants. (A-F) Fold induction of *IFNB1, TNFA, CCL5, IL6, CXCL10* and *MX1*, respectively, by different variants of gRNA including *in vitro* transcribed gRNA (IVT), chemically modified and unmodified full-length synthetic gRNA (FL-MO and FL-UM), duplexed synthetic gRNA chemically modified (Du-MO) and unmodified (Du-UM) in THP1 derived macrophages. Poly(I:C) serves as a control. (G) and (H) The immune inducing capability of gRNA variants were verified in 293T cells and pDC-gen2.2 cells by measuring the fold change of *IFNB1*. (I) Fold induction of *IFNB1* by IVT-gRNA in THP1 derived macrophages using Lipofectamine 2000, RNAi/MAX and Viromer RED. DsDNA (60-mer) serves as a control. All gRNAs target to *CCR5* locus.

### The mechanism of IVT-gRNA induced immune response

The sequence and secondary structure of a *CCR5* locus targeting IVT-gRNA with a 5’ triphosphate (ppp) moiety is illustrated in Fig. 2A. We first investigated the mechanism of IVT-gRNA induced innate immune sensing using 293T cell lines knockout of three cytoplasmic RNA sensors – retinoic acid-inducible gene-I (*RIG-I*), melanoma differentiation-associated gene 5 (*MDA5*) and protein kinase R (*PKR*). While knockout of *MDA5* and *PKR* preserved *IFNB1* induction, knockout of *RIG-I* completely abolished IVT-gRNA-induced *IFNB1* expression indicating RIG-I to be the primary sensor of IVT-gRNA (Fig. 2B). Using the same setting, we investigated mechanism to sense the unmodified Cas9 mRNA and found reduced *IFNB1* induction only when *RIG-I* was knockout (Supplemental_Fig_S3A). Next, we investigated which adaptor molecule for nucleic acids sensors was involved in the IVT-gRNA sensing pathway. We found that cells lacking mitochondrial antiviral-signaling protein (*MAVS*), but not cells lacking stimulator of IFN genes (*STING*) exhibited reduced capacity to evoke *IFNB1* expression following IVT-gRNA transfection (Fig. 2C). RIG-I recognizes the biphosphate and triphosphate moieties in 5’-terminus of double or single stranded RNAs, AU- or polyU/UC-rich region, and specific secondary structure of viral genome (Hornung et al. 2006; Pichlmair et al. 2006; Goubau et al. 2014; Lassig et al. 2015; Sanchez David et al. 2016). Notably, cells knockout of TIR-domain-containing adapter-inducing IFN-beta (*TRIF*) were able to efficiently promote *IFNB1* production but significantly lower than the wild-type counterpart (Fig. 2D and Supplemental_Fig_S3B). To investigate the mechanism in IVT-gRNA sensing, we pre-treated the IVT-gRNA with Alkaline Phosphatase (CIP) to remove the 5’ triphosphate group before transfection. This treatment reduced the induction of *IFNB1* in THP1-derived macrophages (Fig. 2E). Furthermore, the reduced induction of *IFNB1* was recapitulated when capped IVT-gRNA was used for transfection (Fig. 2E). As the IVT-gRNA can be assembled into ribonucleoprotein (RNP) with Cas9 protein *in vitro*, we therefore ask whether preassembly can minimize IVT-gRNA sensing. Indeed, we found significantly lower of *IFNB1* production likely due to preassembly excluded the binding of RIG-I to gRNA (Fig. 2F). Together, these data suggest RIG-I recognition to the 5’ triphosphate group but not the secondary RNA structure is the main contributor to IVT-gRNA induced immune sensing.

**Fig. 2.**
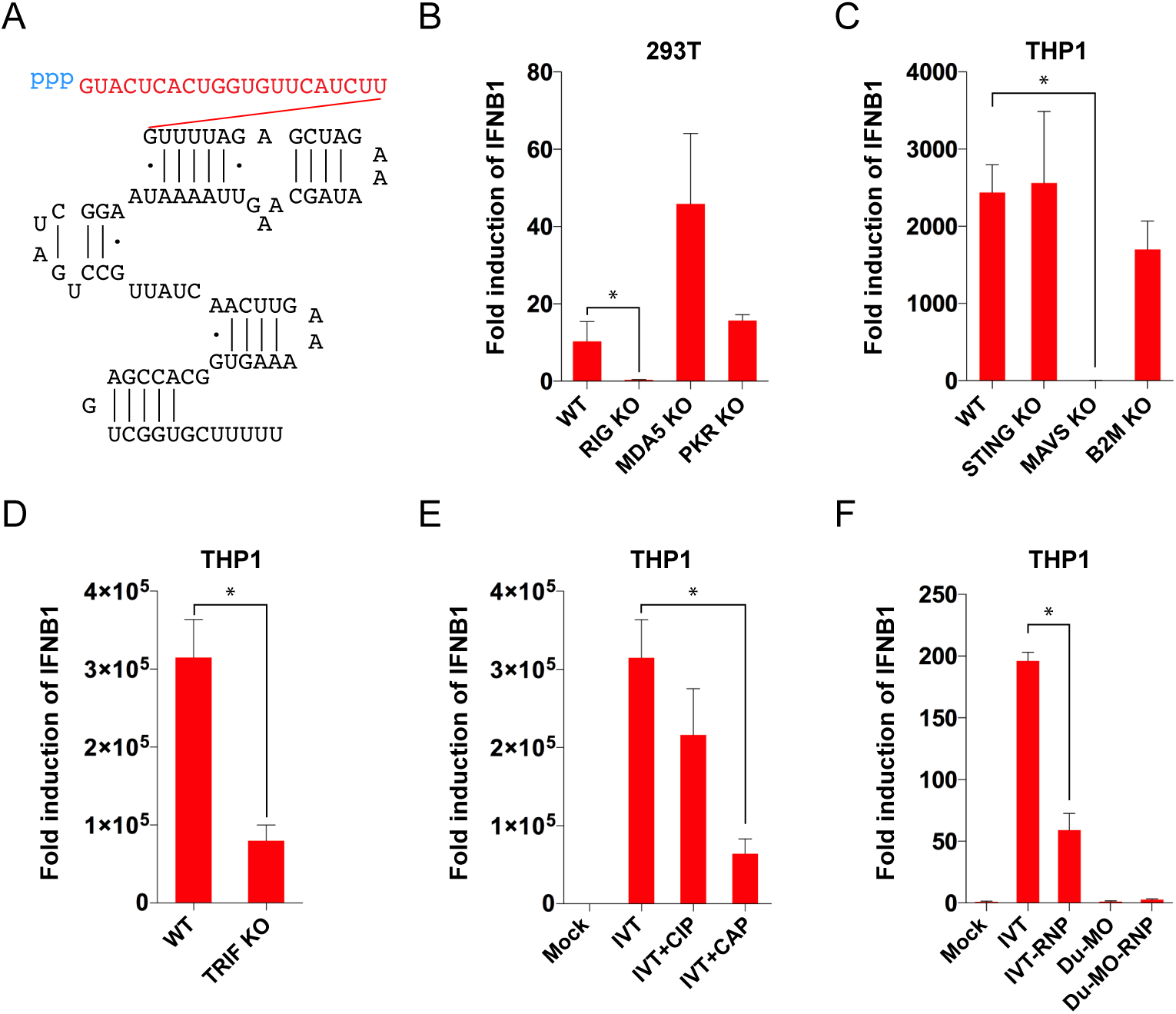
The mechanism of IVT-gRNA induced innate immune response. (A) Sequence and secondary structure of *in vitro* transcribed gRNA targeting to *CCR5* locus. The first three blue letters in lower case indicate triphosphate modification and the followed red letter in upper case is the *CCR5*-locus targeting sequence. The black letter in upper case is backbone of IVT-gRNAs. (B) Evaluation of the innate immune induction by IVT-gRNA in 293T cells knockout of RNA sensor *RIG-I, MDA5* and *PKR*, respectively. (C) and (D) Examining the innate immune induction by IVT-gRNA in THP1 derived macrophages knockout of adaptor protein *STING, MAVS* and *TRIF*, respectively. *B2M* knockout line serves as a control. (E) Fold induction of *IFNB1* by IVT-gRNA, IVT-gRNA pre-treated with CIP and capped IVT-gRNA. (F) Fold induction of *IFNB1* by IVT-gRNA and IVT-gRNA assembled with recombinant Cas9 protein (RNP). The chemically modified duplexed synthetic gRNA and its Cas9 assembled format serve as negative controls.

### Gene disruption efficiency of gRNA variants

To compare the genome-editing efficiency of different *in vitro* transcribed and synthetic gRNAs, we generated a 293T cell line constitutively expressing Cas9, and performed T7 endonuclease I (T7EI) assay following gRNA transfection. All the synthetic gRNAs showed higher efficiency than IVT-gRNA, in particular, the chemically modified gRNAs (Fig. 3A). Although addition of a cap to 5’-end of IVT-gRNA was detrimental for obtaining efficient genome-editing, uncapped IVT-gRNA maintained 10% efficiency (Fig. 3A). When the Cas9 was expressed following transfection of chemically modified Cas9 mRNA which is immune silent, IVT-gRNA, full-length unmodified and full-length modified gRNAs showed efficiency of 10%, 14% and 22%, respectively, similar pattern as in 293T-Cas9 cells (Fig. 3B). However, when we preassembled the gRNAs with Cas9 protein to RNP, IVT-gRNA and full-length unmodified gRNA presented same efficiency (Fig. 3C). Notably, preassembly did not remove the gap of efficiency between full-length unmodified and modified gRNA (Fig. 3C) – a phenomenon that has been noticed previously (Hendel et al. 2015). To investigate influences of RNA sensors on the efficiency of genome editing, we generated *RIG-I, PKR* and *MDA5* knockout cell lines derived from the HEK293T-Cas9 cells. In all tested knockout and wild-type 293T cells, chemically modified full-length gRNA consistently had higher gene disruption efficiency than its unmodified counterpart indicating factors other than the innate RNA-sensing proteins being responsible for lower gene editing by unmodified full-length gRNA (Fig. 3A, 3D-F). In *RIG-I* knockout, but not *PKR* and *MDA5* knockout cell line, IVT-gRNA showed same level of gene disruption efficiency as unmodified full-length gRNA reflecting competition between Cas9 and RIG-I protein to bind to IVT-gRNA (Fig. 3D-F).

**Fig. 3.**
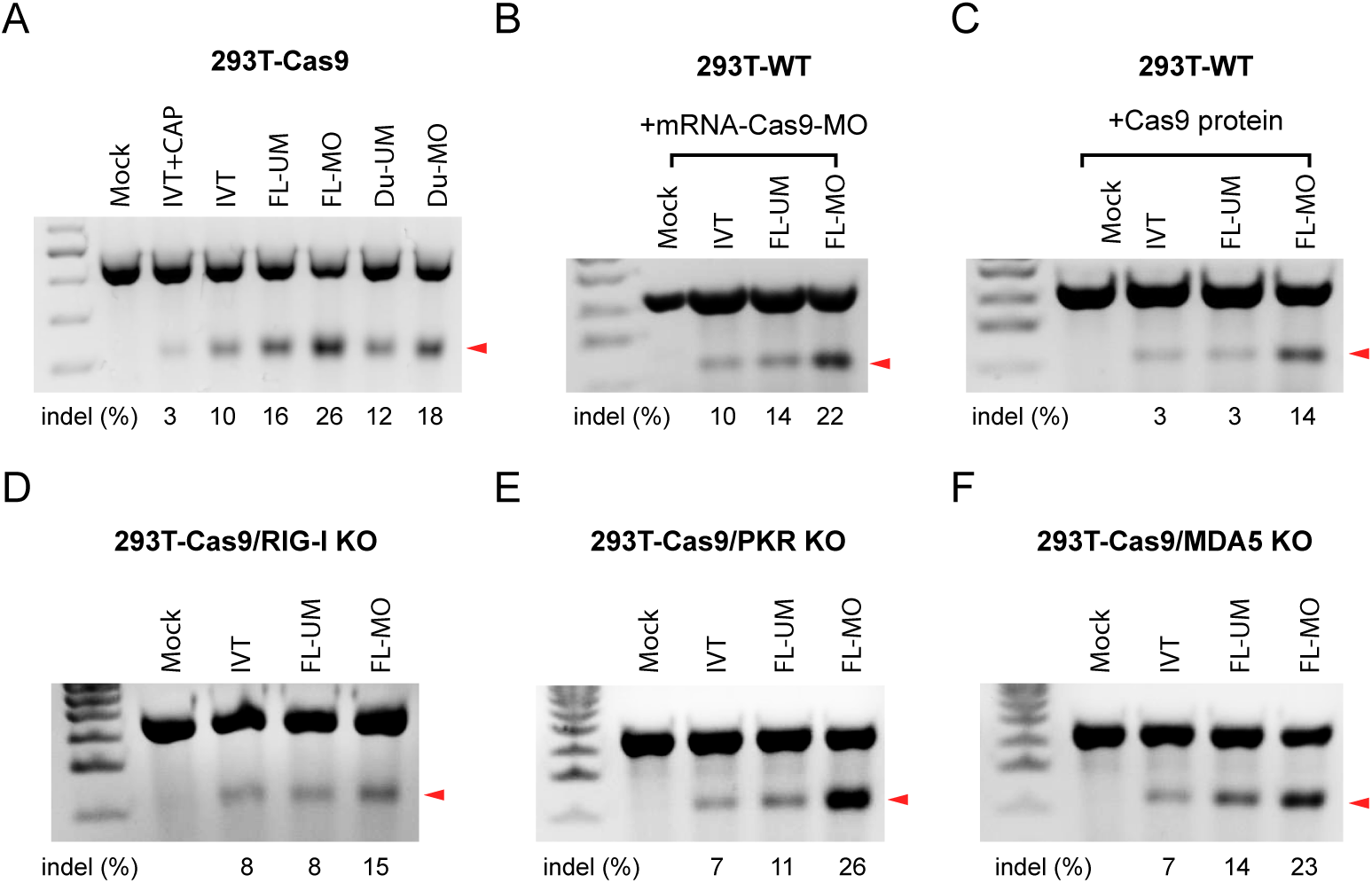
The nuclease activity associated with gRNA variants examined by T7EI assay. (A) Comparison of nuclease activity of a variety of gRNA in a Cas9-expressing 293T line. From left to right, capped *in vitro* transcribed gRNA (IVT+CAP), *in vitro* transcribed gRNA (IVT), synthetic full-length gRNA with (FL-MO) or without chemical modification (FL-UM), synthetic and duplexed gRNA with (Du-MO) or without chemical modification (Du-UM). (B) Nuclease activity of IVT, FL-UM and FL-MO gRNA in wild-type 293T cells. The Cas9 were provided from modified Cas9 mRNA (Cas9-mRNA-MO) which does not elicit innate immune response. (C) Gene disruption induced by gRNA variants preassembled with recombinant Cas9 proteins in 293T WT cells. (D-F) Nuclease activity of IVT, FL-UM and FL-MO gRNA in *RIG-I, PKR* and *MDA5* knockout 293T cell lines. All the knockout lines consistently express Cas9. All gRNAs target to *CCR5* locus.

### The antiviral effects of dual functioning gRNA and the mechanism

Many viruses still cause significant threat to human health. For instance, there is no sterilizing cure for HSV and hepatitis B virus (HBV) infection. CRISPR system has been adapted for treatment of DNA virus (Seeger and Sohn 2014; Seeger and Sohn 2016; van Diemen et al. 2016). So far, the antiviral function of CRISPR has been purely relying on the nuclease activity (Soppe and Lebbink 2017), which makes it less attractive in practice due to the challenge to deliver CRISPR to every infected cell. Innate immune sensing of virus leads to expression and secretion of IFNs which further induce expression of hundreds of IFN-stimulated genes (ISGs) encoding antiviral proteins - thereby protecting both infected and non-infected bystander cells (Paludan 2015). However, viruses have developed mechanisms to escape detection by host cells. For instance, HSV-1 encodes several proteins that antagonize intracellular nucleic acid sensors (Beachboard and Horner 2016; Christensen and Paludan 2017). HSV-1 initiate infection in epithelial cells and subsequently establish latency in sensory neurons. To study the antiviral action of immunostimulatory gRNAs, we chose keratinocytes which are of the epithelial cell linage, and an important target for HSV-1 infection. First, we designed gRNAs targeting HSV-1 and examined whether they induced *IFNB1* expression in HaCaT keratinocytes. We found IVT-gRNA induced robust *IFNB1* expression in HaCaT cells while no *IFNB1* production from the FL-MO counterpart (Fig. 4A) using a HSV-1 *UL8* targeting gRNA sequence of which the nuclease ability had been characterized previously (van Diemen et al. 2016). Interestingly, however, HSV-1 infection in HaCaT did not induce detectable *IFNB1* and *MX1* mRNA nor type I IFN bioactivity (Supplemental_Fig_S4A-C). As unmodified Cas9 mRNA also induces cytokines (Supplemental_Fig_S2), we examined whether we could maximize immune responses by combining IVT-gRNA and unmodified Cas9 mRNA instead of its modified counterpart. Indeed, IVT-gRNA has a tendency to induce more *IFNB1* production when combined with unmodified Cas9 mRNA instead of modified one (Fig. 4B). Notably, the immune maximized combination did not affect the efficiency of gene disruption when testing on endogenous *CCR5* locus (Fig. 4C). Next, we compared the antiviral activity of different CRIPSR combination against HSV1-GFP by measuring the percentage of GFP positive cells. We found the immune silent combination - full-length unmodified gRNAs and modified Cas9 mRNA only reduced GFP modestly but significantly compared to mock-treated control cells (Fig. 4D). In contrast, in the immune maximized group, IVT-gRNA and unmodified Cas9 mRNA strongly suppressed viral gene expression (Fig. 4D). Therefore, we chose the immune maximized combination for further investigation. We evaluated the antiviral activity of two gRNAs targeting to HSV-1 *UL8* and *UL29* gene, respectively. On day 1, both IVT-gRNAs suppressed HSV1-GFP infection significantly (Fig. 4E and Fig. 4F, and Supplemental_Fig_S4D and 4E, left column). On day 2, the virus infection did progress even in the presence of IVT-gRNA (Fig. 4G and Fig. 4H, and Supplemental_Fig_S4E, right column and 4F). However, the IVT gRNA-treated cells preserved intact colony structure in sharp contrast to the mock which failed to form large colonies highlighted by dash lines (Fig. 4H and Supplemental_Fig_S4E, right column) indicating the IVT-gRNAs provide bystander protection to the neighbouring non-transfected cells. In addition, the robust antiviral effects from IVT-gRNA was replicated by the plaque assay using the supernatants from IVT-gRNA treated cells from day 1 (Fig. 4I and Supplemental_Fig_S4G). The bystander *versus* genome-editing-mediated protection from IVT-gRNA can be distinguished by microscopy, since not all cells are transfected with the delivered RNAs. We compared the antiviral activity of immunostimulatory and immune silent RNAs. Interestingly, only the immunostimulatory RNA, but not the immune silent RNAs led to protection of large areas of the infected cell culture (Supplemental_Fig_S4H). To examine whether type I IFNs participated in IVT-gRNA-mediated antiviral responses, we generated a *IFNAR2* knockout HaCaT line which does not respond to type I IFNs (Supplemental_Fig_S5A, B). In this setting, the immunostimulatory RNAs, i.e. IVT-gRNA and unmodified Cas9 mRNA induced relatively strong antiviral activity but to a much less extent than in wild-type cells (Fig. 4J). Importantly, *IFNAR2* KO cells, failed to provide bystander protection to non-transfected cells as no large intact colony was found (Fig. 4K). Collectively, these data demonstrate that the type I IFN system provides potent adding benefits to the CRISPR-mediated antiviral activity.

**Fig. 4.**
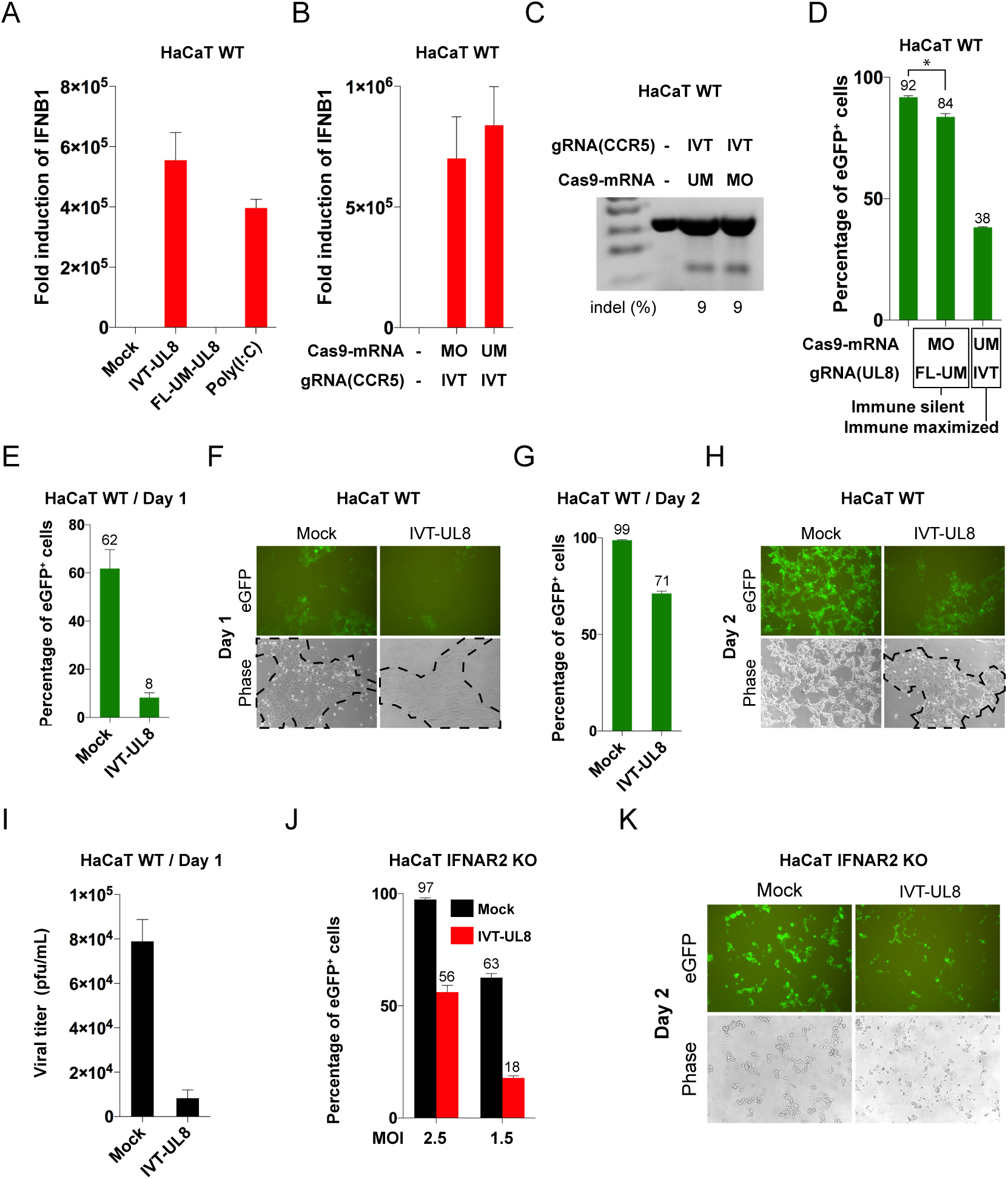
The antiviral effect of IVT-gRNA *in vitro*. (A) Fold induction of *IFNB1* by HSV-1 *UL8* targeting *in vitro* transcribed (IVT-UL8) and full-length unmodified gRNAs (FL-UM-UL8) and Poly(I:C) in HaCaT cells. (B) Fold induction of *IFNB1* by a combination of Cas9-mRNA-MO and IVT-gRNA, and a combination of Cas9-mRNA-UM and IVT-gRNA in HaCaT cells. The gRNA targets to endogenous *CCR5* locus. (C) T7EI assay analysis gene disruption at *CCR5* locus by IVT-gRNA supplemented with unmodified or modified Cas9 mRNA. Percentage of gene disruption (indel) is shown at the bottom. (D) Antiviral effects by ‘immune silencing’ combination (Cas9-mRNA-MO and FL-UM) and ‘immune maximized’ combination (Cas9-mRNA-UM and IVT-gRNA). The gRNA targets to HSV-1 *UL8* gene. (E-I) Antiviral effects of IVT-gRNA in HSV1-GFP infected HaCaT cells. CRISPR was provided by transfection of IVT-gRNA and unmodified Cas9 mRNA 1 h after HSV1-GFP infection (MOI=1.5). The gRNA targets to HSV-1 *UL8*. (E) Flow cytometry analysis the percentage of eGFP^+^ cells 24 h after infection. (F) Representative fluorescent and phase contrast photograph of CRISPR treated cells from (E). (G) Flow cytometry analysis the percentage of eGFP^+^ cells 48 h after infection. (H) Representative fluorescent and phase contrast photograph of CRISPR treated cells from (G). (I) Plaque assay analysis of infectious virus in supernatants harvested from (E). (J) Antiviral effects of IVT-gRNA in HaCaT cells knock out of *IFNAR2* indicated by percentage of eGFP^+^ cells 24 h after HSV1-GFP infection (MOI=1.5 or 2.5). The gRNA targets to HSV-1 *UL8*. The Cas9 was provided by unmodified Cas9 mRNA. HaCaT cells were treated same as (E). (K) Morphology of HaCaT cells knockout of *IFNAR2* two days after HSV1-GFP (MOI=1.5) infection and subsequent IVT-gRNA and unmodified Cas9 mRNA (immune maximized combination) co-treatment.

### The antiviral effects of dual functioning gRNA *in vivo*

To strengthen the concept of using dual functioning IVT-gRNA for antiviral purpose, we adopted a HSV-1 intraperitoneal infection model. As shown Fig. 5A, we added IVT-gRNA 24 h after infection, to mimic a clinically relevant situation with initiation of treatment after infection. Interestingly, intravenous injection of IVT-gRNA encapsulated in nanoparticles, reduced viral load to below detection limit in the spleen in contrast to what was observed in the vehicle treated group (Fig. 5B). In agreement with this, we observed fewer virus-positive cells in the liver of mice receiving IVT-gRNA, which also generally stained less intensely for virus antigen (Fig. 5C). These results suggest that IVT-gRNA possess potent antiviral activity *in vivo*.

**Fig. 5.**
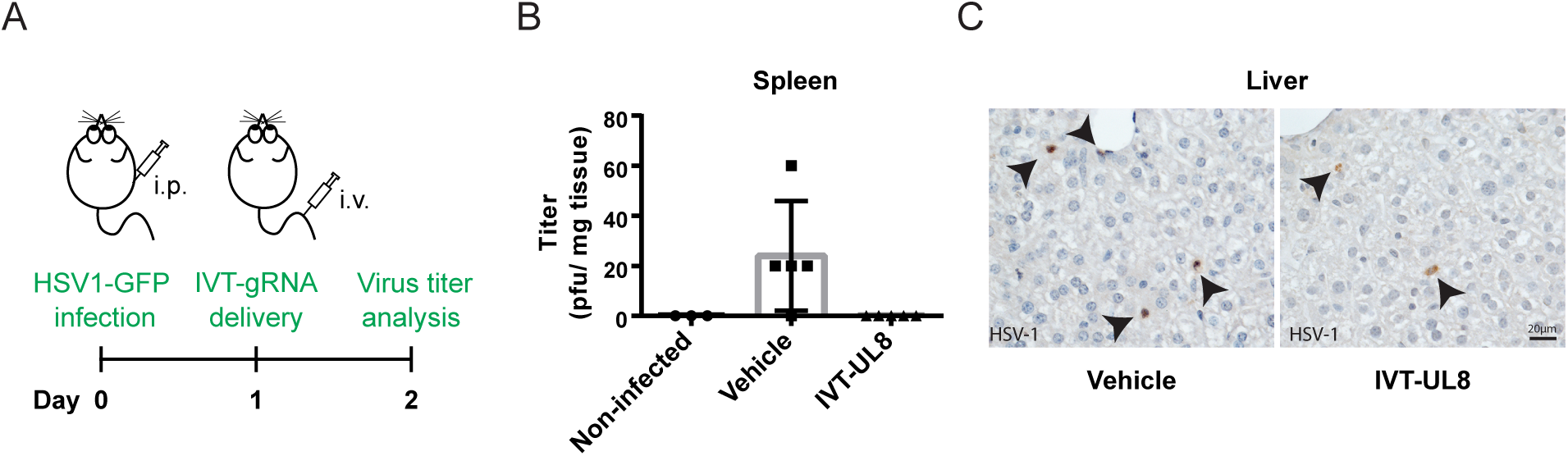
The antiviral effect of IVT-gRNA *in vivo*. A. Treatment scheme. Cas9+ mice were infected i.p. with HSV1-GFP. One day later, the mice were treated with IVT-gRNA encapsulated into nanoparticles and delivered i.v. Spleens and livers were harvested 24 h post infection for further analysis. B. Virus titers in the spleens were determined by plaque assay. C. Liver sections were stained with anti-HSV-1 antibody. Black arrows indicate HSV-1 positive cells. Scale bar = 20 μm.

## Discussion

In a recent publication by Kim et al., the authors investigated innate immune sensing of IVT-gRNA and found that 1) IVT gRNA induces type I IFN, 2) IVT-gRNA is sensed by RIG-I, a *DDX58* encoding protein, 3) Removing IVT gRNA 5′-triphosphate group silences immune response(Kim et al. 2018). While our study confirms the major findings in above-mentioned paper, we provide more detailed investigation regarding mechanism of gRNA sensing. While Kim et al. are interested in avoiding innate immune sensing of IVT-gRNA for gene editing in T cells(Kim et al. 2018), we are interested in exploring the dual function of IVT-gRNA – gene editing and immune stimulating – for antiviral purpose.

Briefly, in this report we have evaluated the immune properties of a variety of commonly used gRNAs and find that only *in vitro* transcribed gRNA induce potent innate immune responses while synthetic gRNAs do not, irrespective of chemical modifications. In agreement with previous reports on *in vitro*-transcribed RNA and siRNAs (Kato et al. 2006; Marques et al. 2006), we show that the RIG-I/MAVS pathway is largely responsible for sensing of IVT-gRNAs. We demonstrate that *in vitro* transcribed gRNA is much more efficient in suppressing HSV-1 infection than synthetic gRNA due to the dual ability to mediate both immune activation and gene disruption. We further demonstrate IVT-gRNA efficiently suppressing HSV-1 *in vivo*. The antiviral activity may be further strengthened by adding 5’ triphosphate to the synthetic chemically modified gRNA to enhance the nuclease activity or using improved delivery methods. Many million people worldwide are persistently infected with viruses. For instance, approximately 400 million people are infected with HBV. The lack of clearance is at least partially contributed by lack of immunological DNA sensing in hepatocytes (but with competent RNA sensing pathway) and failure to induce type I IFN (Sato et al. 2015; Thomsen et al. 2016). Based on the presented current proof-of-concept study, we therefore propose that dual ability IVT-gRNAs have therapeutic potential for antiviral treatment against a range of viruses including HSV-1 and HBV. This urges further exploration.

## Methods

### Mice

The CRISPR/Cas9 knock-in mice constitutively express Cas9 endonuclease (B6J.129(Cg)-Igs2^tm1.1(CAG-cas9*)Mmw^/J) were obtained from The Jackson Laboratory. All experiments were carried out at the University of Aarhus and approved by Danish government authorities. In the infected groups, mice were infected with HSV1-GFP (10^6^ PFU/mouse) at 1-month age by IP injection. The infected mice were either tail vein injected with vehicle (phosphate buffered saline) or IVT-gRNA (40 μg/mouse) complexed with in vivo-jetPEI®-Gal (Polyplus).

### Cells lines

To generate 293T cell line knockout of *PKR*, a gRNA sequence (5’-ttatccatggggaattacat-3’) targeting to *PKR* was cloned into lentiCRISPRv2 puro plasmid (Addgene plasmid # 98290). The acquired gRNA plasmid was then used to produce lentiviral vector for infection of wild-type 293T cells. The puromycin resistant clones was expanded and verified for knockout by Sanger sequencing of multiple bacteria clones. 293T knockout of *RIG-I* (Zhu et al. 2014) and *MDA5* are kind gifts from Veit Hornung (Institute for Clinical Chemistry and Clinical Pharmacology, University of Bonn, Germany). A Cas9 expression cassette was introduced to *RIG-I* and *MDA5* knockouts by infection of a lentiviral vector encoding Cas9 gene. To generate THP1 knockout of *TRIF* which functions in the *TLR3* signaling pathway, we designed a gRNA (5’-gatgaggcccgaaaccggtg-3’) and cloned into the lentiCRISPRv2 puro (Addgene plasmid #98290). Lentiviral vector was then packaged and used to infected THP1 cells. Four days after infection, cells were plated at a dilution sufficient to obtain single cell clones. These clones were expanded into larger cultures where they were validated by Western blot analysis (Supplemental_Fig_S3B). To generate HaCaT knockout of *IFNAR2* which encodes a subunit of the IFNAR heteromer, a guide sequence (5’-caccggtatttcatatgattcgcc-3’) was cloned into the lentiCRISPRv2 puro plasmid (Addgene plasmid # 98290). The HaCaT cells were transfected with the plasmid using the Lipofectamine 2000 (Invitrogen) according to the manufacturer’s protocol. Twenty-four hours post transfection, the cells were seeded in a dilution sufficient to obtain single cell clones after the puromycin selection. The puromycin selection (2 µg/mL) was initiated 48 h post transfection and continued for 72 h. Hereafter, single cell clones were expanded and validated by Sanger sequencing of bacteria clones and functional analysis (Supplemental_Fig_S5A and B).

### Cell cultures and transfection

THP-1 and pDC-gen2.2 cells were maintained in RPMI 1640 medium (Lonza). 293T, HaCaT cells and their knockout cells were maintained in DMEM (Lonza). Both RPMI and DMEM were supplemented with 10% fetal calf serum (FCS, Sigma-Aldrich), 2 mM L-glutamine (Sigma-Aldrich) 100 U/mL penicillin, 100 lg/mL streptomycin (Gibco). To boost cytokine production, THP-1 cells were differentiated into macrophage-like cells with 150 nM phorbol 12-myristate 13-acetate (PMA) (Sigma) before experiment. pDC-gen2.2 cells were cultured on MS-5 irradiated feeder cells. The cells were transfected using Lipofectamine 2000 (ThermoFisher Scientific) for most experiments unless specially indicated where cells were transfected with RNAi/MAX (ThermoFisher Scientific) and Viromer RED (Lipocalyx). Poly(I:C) was purchased from InvivoGen. DsDNA is a 60-mer oligonucleotide derived from the HSV-1 genome.

### RT-qPCR

1.5 × 10^5^ THP-1 derived macrophage-like cells were transfected with different variants of gRNAs or mRNA-Cas9. 6 h after transfection, the cells were harvested for total RNA extraction using the High Pure RNA Isolation kit (Roche). 10^5^ HaCaT cells were infected with HSV1-GFP at a MOI of 2.5 or transfected with 2.5 μg of IVT-gRNA or poly(I:C), and harvested for RNA extraction 16 h later. All the cytokines were measured with the TaqMan RNA-to-Ct 1-step kit (Applied Biosystems) using on the Aria Mx Real Time PCR System, and was normalized to Actβ. The used TaqMan gene expression assays were *IFNB1* (Hs01077958_s1), *TNFA* (Hs00174128_m1), *CCL5* (Hs00174575_m1), IL-6 (Hs00985639_m1), *CXCL10* (Hs01124252_g1), *MX1* (Hs00895608_m1), *CCL2* (Hs00234140_m1), and *ACTB* (Hs00357333_g1).

### HEK-Blue Assay

The level of bioactive type I IFN in supernatants from mock, IVT-gRNA, poly(I:C) or HSV1-GFP treated cells was determined using HEK-BlueTM IFN-alpha/beta cells (Invivogen), according to manufactures instructions, by monitoring the activation of the ISGF3 pathway. HEK-Blue cells has a functioning type I IFN signalling pathway since they are constructed to stably express the human *STAT2* and *IRF9* genes. Furthermore, the cells are designed with the secreted embryonic alkaline phosphatase (SEAP) encoding gene under control of the IFN-alpha/beta inducible promotor from *ISG54*. Thus, upon type I IFN stimulation, the HEK-Blue cells secreted SEAP which is detected using its detection reagent QUANTI-Blue™ by an absorbance shift measured at 620 nm using ELx808 absorbance microplate readers (BioTek).

### T7 endonuclease I assay

293T-Cas9 were transfected with gRNAs alone and 293T cells were co-transfected with gRNAs and Cas9 mRNA or recombinant protein using Lipofectamine 2000 (ThermoFisher Scientific). 48 h after transfection, the cells were harvested and incubated with QuickExtract (Epicentre). The yields were used as templates for PCR amplification using Phusion Master Mix (ThermoFisher Scientific). The PCR products were incubated with 1 µl T7EI nuclease (NEB) in the presence of NEB buffer 2 at 42°C for 30 min after denaturation and re-annealing. The cleavage products were separated on a 1.5% agarose gel and stained with ethidium bromide. The percentage of indels was determined as previous description (Cai et al. 2014; Cai et al. 2016). The primers used to amplify *CCR5* locus are: forward primer 5’-aaaacagtttgcattcatggagggc-3’ and reverse primer 5’-agaagcctataaaatagagccctgt-3’.

### Flow cytometry and fluorescent microscope analysis

HaCaT cells were seeded on 12-well plate at a density of 10^5^/well and subjected to HSV1-GFP infection with MOI=1.5 or 2.5 as indicated. The cells were subsequently transfected with corresponding gRNA and Cas9 mRNA 1 h after infection. The GFP expression and cell morphology were recorded by fluorescence microscopy on 24 h (day 1) or 48 h (day 2) after infection.

### Virus plaque assay

HaCaT cells were seeded on 12-well plate at a density of 10^5^/well and infected with HSV1-GFP at a MOI of 1.5. IVT-gRNA and unmodified Cas9 mRNA were transfected 1 h later. The spleens of mice were homogenized with steel beads (Qiagen) in Tissuelyser (II) (Qiagen) for 3 min at frequency (30 s). The supernatants were harvested for viral titration. Briefly, Vero cells were seeded in 5-cm diameter plates in DMEM supplemented with 5% FCS; on the next day, cells media were replaced with 400 µl 2% FCS DMEM and 100 µl serial dilution of the samples. The dishes were shaken every 10 min for 1 hour before adding 8 mL 2% FCS DMEM containing 2% human immunoglobulin (CSL Behring). The plates were further incubated for 2 days. Cells were subsequently stained with 0.03% methylene blue to allow quantification of plaques.

### gRNA and Cas9 RNA

IVT gRNA in the experiments all *in vitro* transcribed using HiScribe Kit. The synthetic duplex and full-length gRNAs with or without modifications are all purchased from Synthego. The chemically modified versions of synthetic gRNA contain 2’-O-methyl analogs and 3’ phosphorothioate internucleotide linkages at the 5’ and 3’ terminal three bases of the gRNA. The *CCR5* gRNA target to 5’-tactcactggtgttcatctt-3’, *UL8* gRNA targets to 5’-ggggcagccataccgcgtaa-3’ and *UL29* gRNA targets to 5’-gcgagcgtacacgtatccc-3’. Both modified and unmodified Cas9 mRNA was purchased from TriLink. The amount of gRNA used was 2.5 µg per transfection and Cas9 mRNA were 1 µg per transfection. The RNP complex was assembled by incubation of gRNA with recombinant Cas9 protein (IDT) by 1:1 molar ratio at room temperature for 5 min.

### Immunohistochemistry

Liver tissues were fixated in 10% formalin, embedded in paraffin, and cut at 4 µm. The tissues sections submitted to antigen retrieval in a citrate buffer and blocked in 5% BSA. The sections were incubated overnight with primary antibody for HSV2 (1/500 dilution, B0116; Dako) and the stain was visualized with appropriate secondary antibody and HRP. Hematoxylin was used to stain cell nuclei.

### Statistical analysis

Data are presented as mean ± SD in all experiments (n⩾3). Student’s t-tests were performed as indicated (* indicates statistical significance, P<0.05).

## Acknowledgement

We thank Line Reinert, Anja Bille Bohn and Charlotte Christie Petersen to help in collecting the flow cytometry data. The work was supported by a grant from The Lundbeck Foundation (R198-2015-171) and the Novo Nordisk Foundation (NNF16OC0022084).

## Author contributions

Y.C. and S.R.P. conceived the study and designed the experiments; Y.C., A.K., S.J.W., and M.K.T. performed experiments; Y.C., A.K., S.J.W., M.K.T. and S.R.P. analyzed data; Y.C. and S.R.P. wrote the manuscript.

